# PP1β opposes classic PP1 function, inhibiting spine maturation and promoting LTP

**DOI:** 10.1101/2023.01.26.525737

**Authors:** Karl Foley, Cody McKee, Abigail Mayer, Archan Ganguly, Daniel Barnett, Nancy Ward, Yu Zhang, Angus C. Nairn, Houhui Xia

## Abstract

Protein phosphatase 1 (PP1) regulates synaptic plasticity and has been described as a molecular constraint on learning and memory. There are three neuronal isoforms, PP1α, PP1β, and PP1γ, but little is known about their individual functions. PP1α and PP1γ are assumed to mediate the effects of PP1 on learning and memory based on their enrichment at dendritic spines and their preferential binding to neurabin and spinophilin, major PP1 synaptic scaffolding proteins. However, it was recently discovered that human de novo PP1β mutations cause intellectual disability, suggesting an important but ill-defined role for PP1β. In this study, we investigated the functions of each PP1 isoform in hippocampal synaptic physiology using conditional CA1-specific knockout mice. In stark contrast to classic PP1 function, we found that PP1β promotes synaptic plasticity as well as spatial memory. These changes in synaptic plasticity and memory are accompanied by changes in GluA1 phosphorylation, GluN2A levels, and dendritic spine density and morphology, including silent synapse number. These functions of PP1β reveal a previously unidentified signaling pathway regulating spine maturation and plasticity, broadening our understanding of the complex role of PP1 in synaptic physiology.

## Introduction

Protein phosphatase 1 (PP1) is a highly conserved serine (S)-threonine (T) phosphatase that regulates a diverse array of cellular processes. The multifunctional roles of PP1 are mediated by distinct holoenzymes consisting of a PP1 catalytic subunit and one or more regulatory subunits (Bollen et al., 2010; Foley et al., 2021a; Heroes et al., 2013). The PP1 catalytic subunit is encoded by three genes, giving rise to PP1α, PP1β, and PP1γ isoforms which have high sequence similarity but differ in their binding affinity to regulatory proteins. The PP1 catalytic subunits possess several shared binding domains that accommodate substrates of varied physicochemical properties, while the regulatory subunits selectively restrict or promote PP1 activity towards specific substrates. Therefore, PP1 holoenzymes can dephosphorylate a wide range of substrates collectively, while individual holoenzymes maintain high substrate specificity. Further, PP1 isoforms participate in distinct but overlapping pools of PP1 holoenzymes.

In the brain, PP1 regulates synaptic transmission and plasticity (Aow et al., 2015; Foley et al., 2022; Gao et al., 2018; Hu et al., 2006; Hu et al., 2007; Jouvenceau et al., 2006; Morishita et al., 2001) and its partial inhibition improves spatial learning and memory in mice (Genoux et al., 2002). Given the potential therapeutic value of inhibiting PP1 in conditions with learning and memory impairments, identifying the associated PP1 holoenzyme and elucidating its molecular function is of great interest. Significant progress has been made through the identification of PP1 isoforms and regulatory proteins enriched at the synapse. Namely, neurabin (*PPP1R9A*) and spinophilin (*PPP1R9B*) are prominent synaptic scaffolding proteins associated with the actin cytoskeleton that target PP1 to the synapse (Terry-Lorenzo et al., 2002), preferentially binding PP1γ, followed by PP1α, and to a minimal extent PP1β (Bordelon et al., 2005). Selective disruption of the neurabin-PP1 holoenzyme via mutagenesis recapitulates the effect of PP1 partial inhibition on synaptic plasticity (Hu *et al*., 2006; Hu *et al*., 2007; Jouvenceau *et al*., 2006). However, non-selective inhibition of PP1 has different effects on synaptic transmission than selective disruption of neurabin-PP1, suggesting there are other PP1 holoenzymes active at the synapse (Hu *et al*., 2007). Consistent with this notion, all three PP1 isoforms and many additional PP1 regulatory proteins are present at the post-synaptic density (Bayes et al., 2012; Bayes et al., 2011; Foley *et al*., 2021a; Li et al., 2016), although their distinct synaptic functions have not been investigated.

While the majority of studies on PP1 in synaptic plasticity have focused on neurabin and spinophilin, which preferentially bind PP1γ and PP1α, recent evidence suggests PP1β has a significant but ill-defined role in neuronal physiology. Several groups have discovered human de novo mutations in PP1β associated with severe intellectual disability (Gripp et al., 2016; Hamdan et al., 2014; Huckstadt et al., 2021; Lin et al., 2018; Ma et al., 2016; Umeki et al., 2019; Zambrano et al., 2017; Zhou et al., 2020), with none yet identified in PP1α and PP1γ. Further, mRNA expression data suggest PP1β could promote learning and memory: high-performing rodents in the Morris Water Maze have significantly lower levels of PP1γ (Haege et al., 2010) but higher levels of PP1β (Burger et al., 2007) in the hippocampus compared to poor-performing rodents.

In this study, we sought to define the functions of each PP1 isoform in hippocampal synaptic physiology. Using conditional CA1-specific knockout (KO) mice, we found that PP1β has distinct and opposing functions to PP1α and PP1γ in regulating synaptic transmission and plasticity as well as spatial learning and memory. Correspondingly, PP1β KO increases S831 GluA1 phosphorylation, decreases GluN2A levels, and increases dendritic spine density and maturation, including silent synapse number.

## Results

### PP1 isoforms differentially regulate basal synaptic transmission

We have previously shown that genetic KO of PP1γ, but not PP1α, in neural precursor cells (NPCs) leads to a reduction in basal synaptic transmission in the hippocampus (Foley *et al*., 2022). Here, we report that KO of PP1β in NPCs is embryonic lethal, similar to previous findings that global PP1β KO is lethal whereas global PP1α and PP1γ KO mice are viable (Dickinson et al., 2016; International Mouse Phenotyping, 2022). We therefore generated CA1-specific conditional KO mice (cKO) to compare the functions of each PP1 isoform by breeding PP1 isoform floxed mice with a CaMKIIα-Cre transgenic line (T29-1). The T29-1 CaMKIIα-Cre line results in delayed postnatal (∼P20) expression of Cre in CA1(Tsien et al., 1996), obviating potential developmental effects of gene knockout. We confirmed CA1-specific Cre expression at 1-1.5 months of age through breeding with an Ai9 reporter line (Figure 1A). Interestingly, cKO of each PP1 isoform had a distinct effect on hippocampal synaptic transmission, as determined by recording field excitatory postsynaptic potentials (fEPSPs) from CA3 Schaffer collateral-CA1 (Sch-CA1) synapses in acute hippocampal slices. PP1γ cKO reduced synaptic transmission whereas PP1α cKO had no effect (Figure 1B-C), consistent with the findings in NPC-KO mice (Foley *et al*., 2022) and the effect of disrupting neurabin-PP1 interaction (Hu *et al*., 2007). To test if an effect of PP1α cKO is masked by PP1γ compensation, we generated a PP1α/γ double cKO mouse. PP1α/γ cKO reduced synaptic transmission (Figure 1D) to a greater degree than PP1γ cKO alone (Figure 1F), suggesting that PP1α plays a secondary, redundant role to PP1γ in regulating synaptic transmission, rather than no role at all. Surprisingly, PP1β cKO increased synaptic transmission (Figure 1E), an unprecedented finding in PP1 synaptic function. Overall, the findings suggest that PP1 isoforms play opposing roles in regulating hippocampal synaptic transmission, with PP1γ and PP1α promoting transmission and PP1β attenuating transmission.

**Fig. 1.**
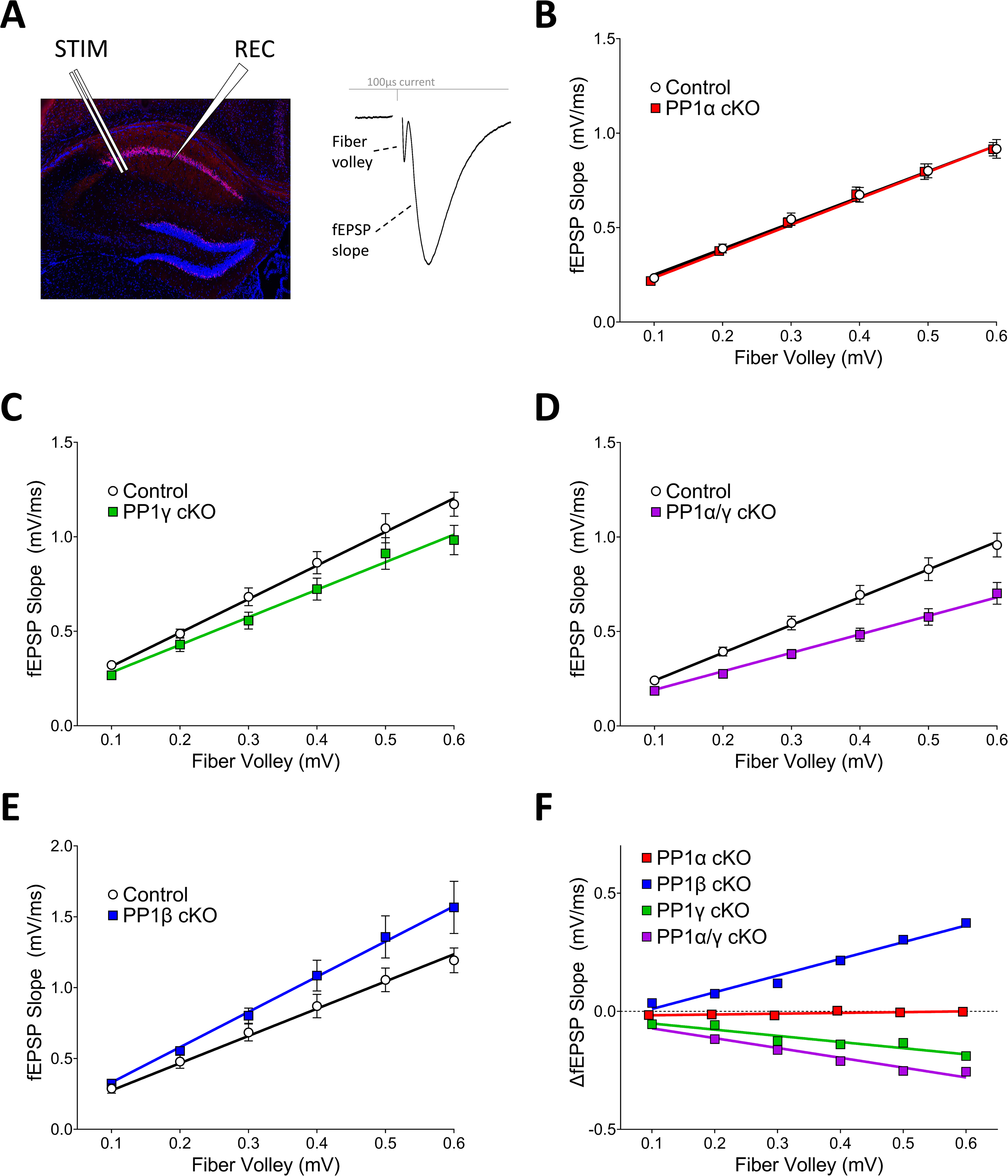
PP1 isoforms play opposing roles in regulating hippocampal synaptic transmission. (**B-E**) (**A**) CA1-specific Cre expression shown by breeding with Ai9 mice, which express tdTomato following Cre-recombination. Field recordings of Sch-CA1 synapses were conducted to assess input-output (IO) curves. (**B)** PP1α cKO does not significantly alter synaptic transmission compared to control littermates (two-way ANOVA, genotype: p=0.6504; Regression: Extra sum-of-squares F-test (F-test), slope: p=0.7769; N=4 (control) and 3 (cKO) mice, n=9 and 10 slices), whereas (**C**) PP1γ cKO significantly decreases transmission (two-way ANOVA, genotype: p<0.01; F-test, slope: p<0.001; N=5 and 3, n=9 and 8) as does (**D**) PP1α/γ cKO (two-way ANOVA, genotype: p=<0.0001; F-test, slope: p<0.0001; N=9 and 10, n=20 and 15). On the other hand, (E) PP1β cKO significantly increases transmission (two-way ANOVA, genotype: p<0.01; F-test, slope: p<0.0001; N=4 and 4, n=11 and 14). (**F**) Comparison of changes in IO curves for PP1 isoform cKO mice normalized to control littermates. PP1α/γ cKO decreases transmission to a greater degree than PP1γ cKO (two-way ANOVA, genotype: p<0.001; F-test, slope: p<0.0001).

### PP1 isoforms differentially regulate LTP

To investigate the roles of PP1 isoforms in synaptic plasticity, we induced LTP via a single bout of high-frequency stimulation. Given the effects on basal synaptic transmission (Figure 1), we chose to focus on PP1α/γ cKO and PP1β cKO mice. PP1α/γ cKO had no effect on classic LTP induction (Figure 2A; 100 pulses, 100Hz) at Sch-CA1 synapses, but significantly increased the potentiation following sub-threshold LTP stimulation (Figure 2B; 25 pulses, 100Hz). In contrast, PP1β cKO significantly attenuated LTP (Figure 2C; 100 pulses, 100Hz). Similar to the effects observed for basal synaptic transmission, the findings suggest PP1β opposes PP1α/PP1γ function in regulating hippocampal synaptic plasticity.

**Fig. 2.**
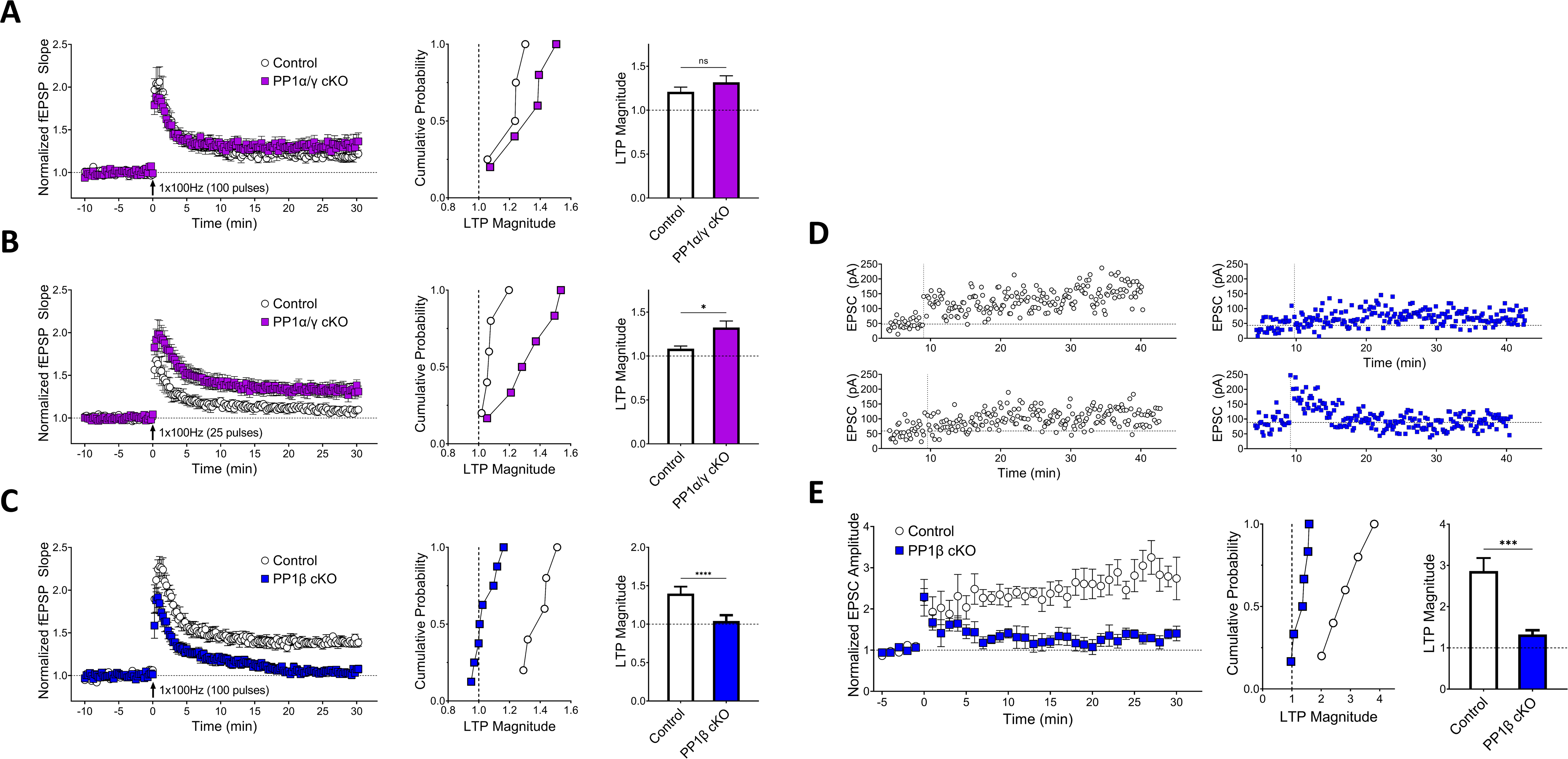
PP1 isoforms play opposing roles in regulating hippocampal synaptic plasticity. (**A**) PP1α/γ cKO does not alter LTP induced by 100 pulses at 100 Hz stimulation (two-tailed t-test: p=0.30, N=3 and 4, n=4 and 5). (**B**) PP1α/γ cKO promotes sub-threshold LTP induced by 25 pulses at 100 Hz stimulation (two-tailed t-test: p<0.05, N=3 and 3, n=5 and 6). (**C**) PP1β cKO attenuates LTP induced by 100 pulses at 100 Hz stimulation (two-tailed t-test: p<0.001, N=3 (control) and 4 (cKO) mice, n=5 and 8 slices). (**D**) Examples of whole-cell recording LTP experiments from control (left) and PP1β cKO (right). LTP was delivered by pairing 100 pulses at 100 Hz at 0 mV holding potential (x-axis dotted line) before 10 min of recording to avoid washout. Y-axis dotted line denotes average baseline response. (**E**) PP1β cKO attenuates LTP induced via the pairing protocol (two-tailed t-test: p<0.001, N=3 (control) and 5 (cKO) mice, n=5 and 6 cells). LTP magnitude was measured as the average fEPSP or EPSC 25-30 min after LTP induction normalized to the baseline.

Many different factors can impair postsynaptic depolarization and thus impair NMDAR activity during LTP stimulation (Granger and Nicoll, 2014). To exclude impaired postsynaptic depolarization as a cause of the LTP deficit in PP1β cKO mice, we induced LTP in CA1 pyramidal neurons using whole-cell patch clamp via the pairing protocol (Figure 2D). In this approach, postsynaptic depolarization is controlled by holding the cell at 0 mV during LTP stimulation. Further, we inhibited GABA_A_ receptors by including 50μM picrotoxin in the bath to exclude altered inhibitory signaling as a mechanism of the LTP deficit. Consistent with the LTP deficit observed via field recordings, we found that PP1β cKO also significantly attenuated LTP induced via the pairing protocol (Figure 2E).

### PP1β cKO impairs spatial learning and memory

Changes in hippocampal LTP are often associated with changes in spatial learning and memory. We therefore sought to test CA1-dependent learning and memory with the allocentric object-place assay (Figure 3A), a variation of novel-object recognition (Langston and Wood, 2009). PP1β cKO mice explored the novel object significantly less than their control littermates (Figure 3B), suggesting impaired short-term spatial memory. PP1α/γ cKO mice did not demonstrate any change in novel object recognition, exploring the novel object to the same degree as their littermate controls (Figure 3F). Neither PP1β cKO nor PP1α/γ cKO mice demonstrated any alteration in the total time exploring both novel and familiar objects (Figure 3C, G). Additionally, there was no change in locomotor activity or anxiety, as measured in open field testing (Figure 3D-E, H-I).

**Fig. 3.**
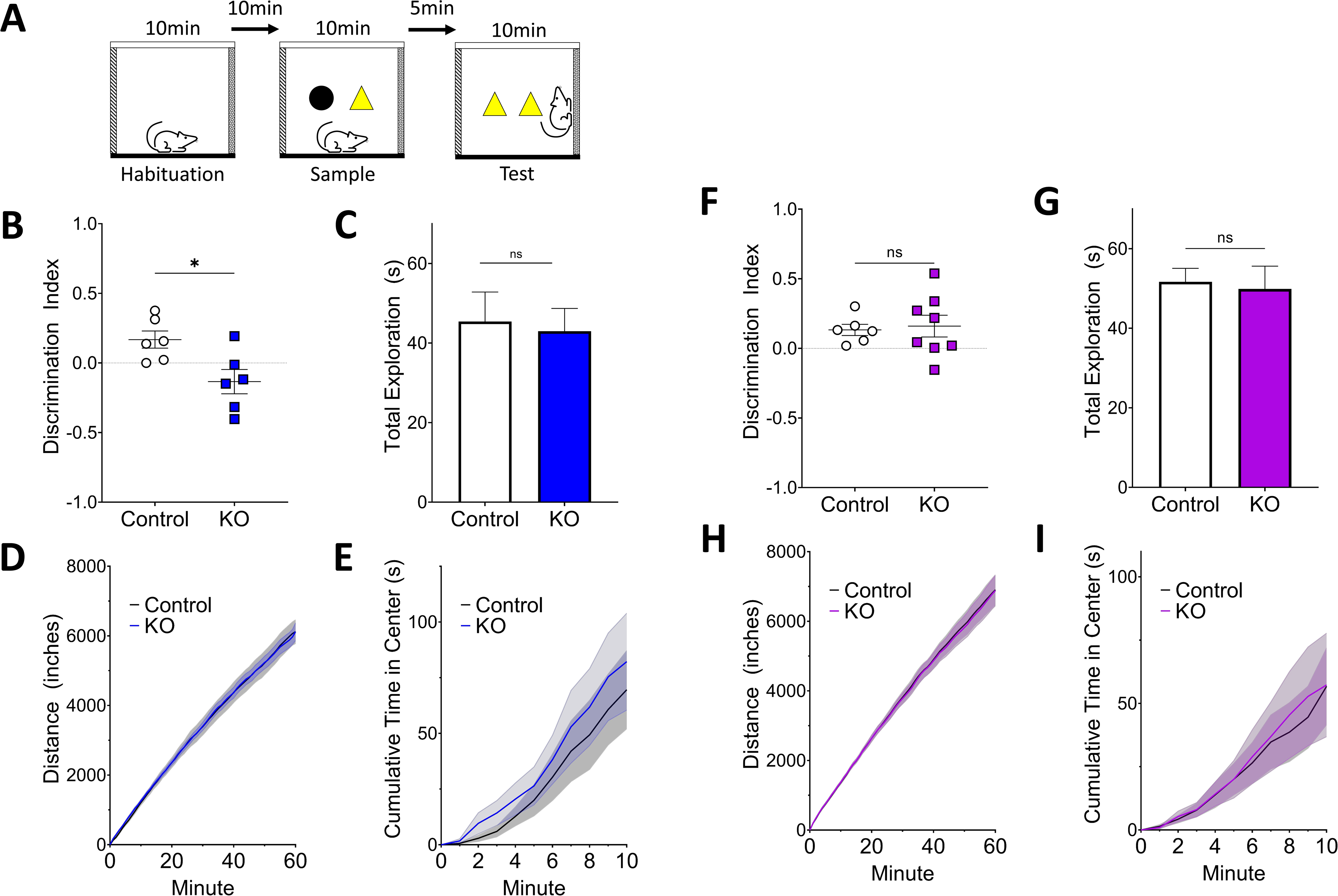
PP1β cKO impairs spatial learning and memory. (**A**) Overview of the allocentric place-object task, a novel object recognition variant sensitive to hippocampal deficits. In the test phase, the novelty of the object is based on its location in the arena. (**B**) PP1β cKO are impaired in novel object recognition compared to their control littermates (two-tailed t-test: p<0.05, N=5 (control) and 6 (cKO) mice, 3 litters). There was no difference in (**C**) total time exploring objects between groups (two-tailed t-test: p=0.98, N=5 and 6). (**D**) Locomotor activity (two-tailed t-test: p=0.82, N=8 and 7), or (**E**) anxiety (two-tailed t-test: p=0.66, N=8 and 7), as measured by time spent in the center during an open field test. (**F**) There was no difference in novel object recognition between PP1α/γ cKO and control littermates (two-tailed t-test: p=0.79, N=6 and 8, 4 litters). PP1α/γ cKO mice showed no difference in (**G**) total time exploring objects between groups (two-tailed t-test: p=0.81, N=6 and 8), (**H**) locomotor activity (two-tailed t-test: p=0.97, N=8 and 10), or (**I**) anxiety (two-tailed t-test: p=0.98, N=8 and 10). Locomotor activity was measured as cumulative distance traveled at 1 hour. Anxiety was measured by time spent in the center of the open field arena over 10 minutes. The standard error of the mean is displayed as error bars or shaded lines.

### PP1β regulates AMPAR phosphorylation and GluN2A levels

Our findings thus far demonstrate that PP1β has distinct, opposing functions to PP1α and PP1γ in the CNS. We therefore sought to identify altered signaling pathways and measure glutamatergic receptor levels. In immunoblotting experiments on CA1 microdissections, we observed no change in several pathways involved in synaptic plasticity and previously shown to be regulated by PP1, including pCREB, pCaMKII, pERK (Figure 4A). However, PP1β cKO mice had significantly increased levels of phosphorylated GluA1 AMPAR at S831 (Figure 4B), a site involved in LTP (Lee et al., 2000; Mammen et al., 1997) that increases AMPAR channel conductance (Derkach et al., 1999) and contributes to GluA1 targeting to the post-synaptic density (PSD) (Diering et al., 2016; Diering and Huganir, 2018) (but see (Hayashi et al., 2000)). There was no significant change in phosphorylation at GluA1 S845 or GluA2 S880, which are dephosphorylated during LTD (Hu *et al*., 2007; Lee et al., 1998), and no change in total GluA1, GluA2, or GluA3 levels (Figure 4B). We also observed a decrease in GluN2A levels in PP1β cKO mice (Figure 4C). Meanwhile, there was no change in GluN2B S1480 phosphorylation, a known PP1 target that regulates GluN2B lateral diffusion (Chiu et al., 2019), or total GluN2B or GluN1 levels (Figure 4C).

**Fig. 4.**
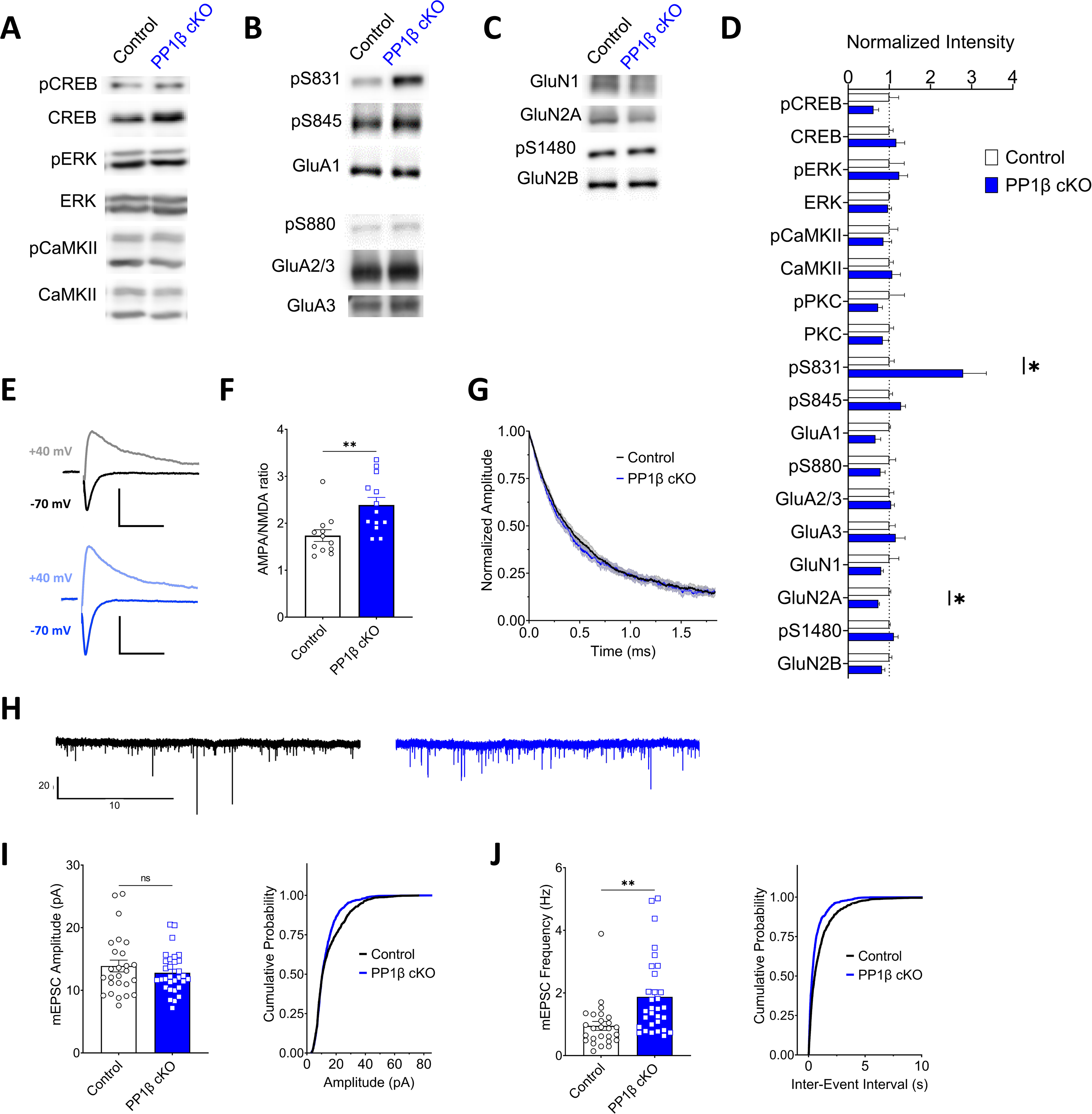
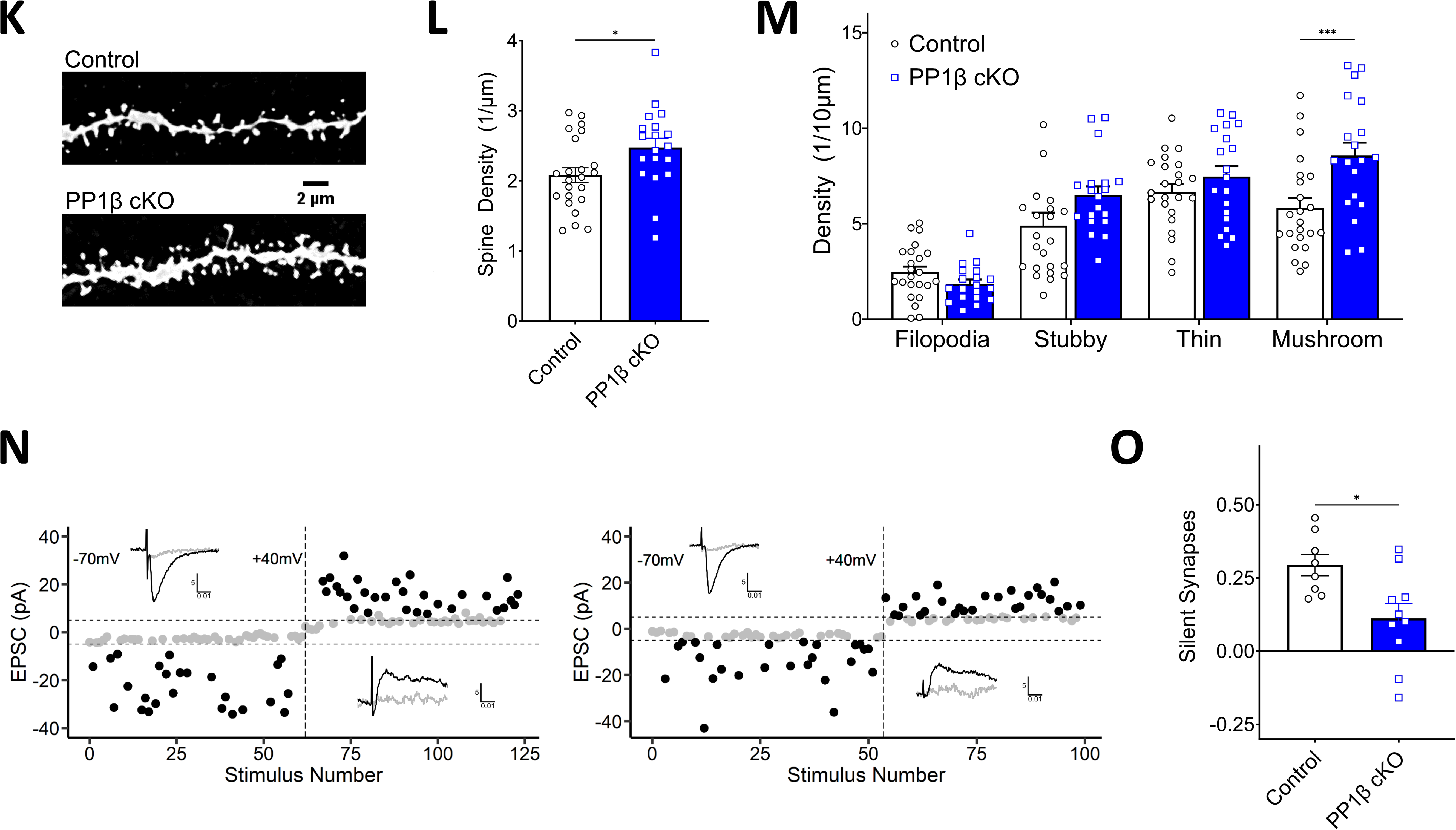
PP1β cKO increases GluA1 phosphorylation, spine density, and spine maturation. (**A-D**) Western blot results from CA1 microdissections of PP1β cKO mice and littermate controls. 3 mice per group. Phosphorylation was normalized to total protein, and total protein was normalized to β-tubulin. (**A**) Signaling pathways relevant to synaptic plasticity. No significant differences were observed. (**B**) AMPAR phosphorylation and total protein levels. S831 GluA1 phosphorylation was significantly increased in PP1β cKO (two-tailed t-test: p<0.05). (**C**) NMDAR phosphorylation and total protein levels. Total GluN2A levels were significantly decreased (two-tailed t-test: p<0.01). (**D**) Quantification of comparisons. Following normalization to total protein or β-tubulin, intensities were normalized to control samples. (**E-G**) Results from AMPAR/NMDAR ratio experiments from CA1 pyramidal neurons. (**E**) Example averaged traces from individual control (top) and PP1β cKO (bottom) neurons held at -70 mV and +40 mV. Stimulation artifacts removed for clarity. Scale bars denote 50 pA and 50 ms. (**F**) PP1β cKO significantly increases the AMPAR/NMDAR ratio (two-tailed t-test: p<0.01, N=3 and 3, n=12 and 13), (**G**) with no effect on the NMDAR decay rate. (**H-J**) Results from mEPSC recordings from CA1 pyramidal neurons. (H) Example mEPSC traces from control (left) and a PP1β cKO (right). (**I**) PP1β cKO has no effect on mean mEPSC amplitude (two-tailed t-test: p=0.31, N=7 (control) and 6 (cKO) mice, n=26 and 31) but alters the amplitude distribution (Kolmogorov-Smirnov: p<0.0001, 1000 randomly sampled events per group). (**J**) PP1β cKO significantly increases mEPSC frequency (two-tailed t-test: p<0.01, N=7 and 6, n=26 and 31) and correspondingly shifts the frequency distribution (Kolmogorov-Smirnov: p<0.0001, 1000 randomly sampled events per group). (**K-M**) Spine number and morphology of CA1 pyramidal neurons. (**K**) Examples of dendritic spines in biocytin-filled CA1 pyramidal cells. (**L**) PP1β cKO significantly increases spine density (two-tailed t-test: p<0.05, N=4 (control) and 3 (cKO) mice, n=23 and 19 dendrites). **(M)** PP1β cKO alters spine morphology (two-way ANOVA, genotype: p<0.05), significantly increasing mushroom spines (Šidák, p<0.001), as categorized by Imaris reconstruction. (**N**) Examples of minimal stimulation experiments from control (left) and PP1β cKO (right) mice. Failures are shown in grey, successful responses in black. Y-axis dotted lines denote 5 pA thresholds used for initial classification. X-axis dotted line denotes the switch to +40 mV. Insets show waveforms from averaged responses at -70 mV (top left) and +40 mV (bottom right); scale bars 5 pA, 10 ms. All events were manually reviewed. (**O**) PP1β cKO decreases the proportion of silent synapses in CA1 pyramidal cells (two-tailed t-test: p<0.05, N=4 and 7, n=8 and 10 cells).

To investigate whether these changes in GluA1 and GluN2A are expressed at the synapse, we measured the AMPAR/NMDAR ratio via whole-cell recordings (Figure 4E-G). As expected from the increase in synaptic transmission and the increase in GluA1 S831 phosphorylation (pS831), we found an increase in the AMPAR/NMDAR ratio in PP1β cKO mice (Figure 4F). However, we observed no change in the decay rate of the NMDAR current (Figure 4G), which would be expected with a change in the relative abundance of synaptic GluN2A and GluN2B subunits. This suggests that GluA1 pS831 phosphorylation but not GluN2A levels are altered at the level of the synapse. Correspondingly, using a non-selective, inducible KO line (iKO; Thy1-CreER) to generate sufficient material for fractionation and immunoblotting of S831 phosphorylation, we found that the increase in GluA1 pS831 was present in the synaptosomal fraction of the hippocampus of PP1β iKO mice (Figure S1A). On the other hand, we found no difference in GluA1 pS831 in PP1α/γ cKO mice (Figure S1B). Further, we did not observe a compensatory increase of the remaining PP1 isoform(s) in PP1β cKO or PP1α/γ cKO mice (Figure S1C-D), suggesting that GluA1 S831 is a unique target of PP1β.

### PP1β regulates spine density, morphology, and silent synapses

Given the increase in synaptic transmission and GluA1 pS831 in PP1β cKO mice, we assessed miniature excitatory postsynaptic currents (mEPSCs) in CA1 pyramidal cells to measure synaptic AMPA receptor (AMPAR) content. Surprisingly, PP1β cKO mice showed a robust increase in mEPSC frequency but not amplitude (Figure 4H-J). Interestingly, although the mean amplitude was not significantly different, there was a significant shift in the cumulative frequency, with a higher proportion of events in the 15-50 pA range in PP1β cKO cells (Figure 4H-I). A change in mEPSC frequency can reflect an increase in the docked vesicle pool or an increase in synapse number. To test for a presynaptic cause, we measured paired-pulse facilitation (PPF) at Sch-CA1 and observed no difference between PP1β cKO and control littermates (Figure S2A). We also observed no difference between slice preparation protocols (Figure S2B-E).

The lack of change in PPF combined with the postsynaptic KO approach strongly suggested an increase in synapse number. We therefore analyzed dendritic spine density in proximal dendrites of CA1 pyramidal neurons by confocal fluorescence microscopy of biocytin-filled neurons (Figure 4K-M). PP1β cKO mice had a significant increase in spine density (Figure 4L), as assessed by Imaris reconstruction of imaged dendrites. The increase in spine density was driven by an increase in mature mushroom spines (Figure 4M).

In addition to morphology, spines can be classified as functional or silent based on their expression or lack of expression of AMPARs, respectively. The specific increase in mature spines in PP1β cKO mice suggests that the increased mEPSC frequency could result from an increase in the number of functional, AMPAR-containing spines in addition to the increase in total number of spines. To test this possibility, we measured AMPAR- and NMDAR-mediated currents in response to minimal stimulation (Figure 4N-O). Stimulation intensity was lowered until approximately 50% of stimulations failed to elicit a response at -70 mV holding potential. We then measured the failure rate at +40 mV to determine the proportion of silent synapses, which exhibit a detectable response at +40 mV but not at -70 mV. PP1β cKO mice showed fewer silent synapses (Figure 4O), suggesting that PP1β cKO increases mEPSC frequency by increasing both the total number and AMPAR-expressing number of synapses.

## Discussion

PP1 is classically viewed as a molecule of forgetfulness, with its inhibition improving spatial learning and memory and promoting LTP over LTD (Genoux *et al*., 2002; Jouvenceau *et al*., 2006). However, PP1 can have broad functions, given that PP1 refers to three isoforms and hundreds of PP1 holoenzymes exist with presumed distinct functions. Here, we demonstrate that PP1β overall opposes PP1γ and PP1α function in synaptic transmission, plasticity, and spatial learning and memory. Whereas PP1α/PP1γ cKO decreases synaptic transmission in Sch-CA1 synapses, consistent with neurabin-PP1 function, PP1β cKO increases synaptic transmission. Similarly, while PP1α/PP1γ cKO promotes LTP, PP1β cKO attenuates LTP. Finally, PP1β cKO impairs spatial learning and memory in a novel object recognition assay, whereas PP1α/PP1γ cKO has no effect. These effects are accompanied by changes in spine morphology, S831 GluA1 phosphorylation, and GluN2A levels. These novel functions of PP1β reveal a previously unrecognized complexity of PP1 function in synaptic physiology.

The distinct roles of PP1β help resolve discrepancies between the effects of non-selective PP1 inhibition and disruption of specific holoenzymes. We previously showed that non-selective PP1 inhibition has no effect on synaptic transmission whereas disruption of neurabin-PP1 decreases synaptic transmission (Hu *et al*., 2007), suggesting the existence of an additional PP1 holoenzyme that opposes neurabin-PP1 function. Our findings suggest that a PP1β holoenzyme mediates this opposing role.

The impairment in spatial learning in PP1β cKO mice stands in stark contrast to the improvement seen with partial PP1 inhibition (Genoux *et al*., 2002). One explanation is that the relevant PP1β holoenzyme remains sufficiently active in the partial inhibition paradigm. In the aforementioned studies, partial inhibition of PP1 was achieved via overexpression of a constitutively active inhibitor-1 fragment (I-1*) (Genoux *et al*., 2002; Jouvenceau *et al*., 2006). It is unclear if all PP1 holoenzymes are equally and proportionally inhibited by I-1*, or if the partial inhibition primarily affects a subset of holoenzymes. I-1-PP1 interaction involves the RVXF binding domain and is predicted to involve the PP1 acidic groove (Liang et al., 2018; Terrak et al., 2004). I-1* may therefore have higher binding affinity to PP1 holoenzymes in which the acidic groove is fully exposed, such as neurabin-PP1 (Ragusa et al., 2010). Phosphoproteomic analysis of the I-1* transgenic mouse would likely resolve these questions and provide valuable insights into the mechanisms by which PP1 partial inhibition improves learning and memory.

The mechanism by which PP1β cKO alters synaptic transmission and plasticity appears to be explained by increased phosphorylation of S831 GluA1. While decreased GluN2A levels may also contribute, our electrophysiology studies did not reveal evidence of decreased GluN2A at the synapse. Phosphorylation of GluA1 at S831 increases AMPAR channel conductance (Derkach *et al*., 1999) and contributes to GluA1 targeting to the PSD under basal conditions (Diering *et al*., 2016; Diering and Huganir, 2018), consistent with the observed increase in synaptic transmission and the shift in the distribution of mEPSC amplitudes. Increased targeting of GluA1 to the PSD via S831 phosphorylation may help account for the decrease in silent synapses in PP1β cKO mice, though the contribution of GluA1 phosphorylation to unsilencing has not been directly investigated. Further, phosphomimetic S831 LTP is accompanied by an increase in GluA1 S831 phosphorylation (Lee *et al*., 2000; Mammen *et al*., 1997), suggesting the increase in pS831 in PP1β cKO may also contribute to the LTP deficit by saturating the contribution of pS831 to potentiation. However, dual-phosphomimetic (S831D/S845D) GluA1 mice show a decrease in the LTP threshold (Makino et al., 2011). On the other hand, the effect of S831D and S845D alone on LTP has not been reported, therefore it is unknown if one or both phosphorylation sites mediates the decreased LTP threshold.

The decrease in total GluN2A levels in PP1β cKO is consistent with the impairment of LTP and spatial learning and memory (Kiyama et al., 1998; Sakimura et al., 1995). However, the decrease in GluN2A was not evident in the decay rate of the combined AMPAR-NMDAR waveform, suggesting the decrease might not localize to the synapse. While the relative contributions of GluN2A and GluN2B to LTP are still debated, classic pharmacologic and genetic ablation studies have demonstrated that GluN2A contributes significantly to LTP at Sch-CA1 (Shipton and Paulsen, 2014). Interestingly, GluN2B-containing NMDARs are enriched in small spines, and GluN2A-containing NMDARs in larger spines (Sobczyk et al., 2005), opening the possibility that, if GluN2A contributes to the observed potentiation deficit, the deficit could vary between spine subtypes.

How does PP1β regulate GluA1 S831 phosphorylation and GluN2A levels? Each function of PP1 is mediated by a specific holoenzyme, consisting of the PP1 catalytic subunit and regulatory protein. It is currently unclear which PP1β holoenzyme, or holoenzymes, mediates the synaptic functions defined in this study, as the neuronal regulatory proteins that bind PP1β specifically have not been characterized. PP1β may directly dephosphorylate GluA1, or act on other relevant regulatory proteins. For example, PP1β cKO may increase GluA1 S831 phosphorylation by promoting CaMKII and PKC activity. We did not observe a change in CaMKII T286 phosphorylation in PP1β cKO mice, but we cannot exclude the possibility that other phosphorylation sites are affected. Similarly, PP1β may directly dephosphorylate GluN2A, a GluN2A scaffolding protein, or another regulatory protein. Several kinases can affect GluN2A surface expression and recycling, including GSK3β, DYRK1A, PKC, CaMKII, and CDK5, either through direct action on GluN2A or its scaffolding proteins (Gardoni et al., 2001; Gardoni et al., 2003; Grau et al., 2014; Li et al., 2001; Mauceri et al., 2007; Monaco et al., 2018; Mota Vieira et al., 2020; Yong et al., 2021). If the PP1β cKO impairments are mediated by unopposed activity of a kinase, it may be possible to rescue the deficits via inhibition of the corresponding kinase.

Although the identities and functions of PP1β holoenzymes in the central nervous system are largely unstudied, MYPT1-PP1β is an attractive candidate holoenzyme. MYPT1-PP1β, also known as myosin phosphatase, regulates the activity of myosin II proteins, including non-muscle myosin II, via inhibitory dephosphorylation of the myosin light chain (MLC) regulatory subunit (Ito et al., 2004). MLC has been implicated in spine maturation, LTP, as well as NMDAR trafficking (Amparan et al., 2005; Bajaj et al., 2009; Hodges et al., 2011; Rex et al., 2010). Given that PP1β opposes classic PP1 function, identifying neuronal PP1β holoenzymes and elucidating their functions presents an exciting avenue for further research.

Human de novo mutations in PP1β cause intellectual disability (Gripp *et al*., 2016; Hamdan *et al*., 2014; Huckstadt *et al*., 2021; Lin *et al*., 2018; Ma *et al*., 2016; Umeki *et al*., 2019; Zambrano *et al*., 2017; Zhou *et al*., 2020). The clinical profile of patients with these mutations resembles a Rasopathy, specifically Noonan syndrome with loose anagen hair. Consistent with this classification, PP1β mutations have been reported to promote Ras signaling (Young et al., 2018). We did not observe a change in pERK, which is downstream of Ras signaling, suggesting PP1β activity may not be important for basal Ras-MEK-ERK signaling and supporting the notion that PP1β mutations represent a gain-of-function. On the other hand, PP1β may be important for activity-induced Ras-MEK-ERK signaling, presenting another means by which PP1β cKO could impair LTP. Further investigation of the neuronal functions of PP1β will improve our understanding of the pathophysiology of these mutations and help generate potential therapeutic targets.

### Limitations of the study

The distinct functions of PP1β identified in this study present exciting opportunities for further delineation of underlying mechanisms. For example, the increase in synaptic transmission along with the decrease in silent synapses could in part be explained by an increase in Ca2+-permeable GluA1 homomeric AMPAR (CP-AMPARs). CP-AMPARs have high channel conductance and are transiently recruited in synapse unsilencing (Morita et al., 2014; Purkey and Dell’Acqua, 2020), Therefore, in addition to GluA1 S831 phosphorylation, the regulation of CP-AMPARs may present another target of PP1β.

While we show in this study that PP1β cKO results in a deficit in spatial learning and memory, our model and behavioral assay are different than the classic study of PP1 function in learning and memory (Genoux *et al*., 2002). Firstly, we utilize a CA1-specific KO instead of a widespread neuronal KO, since global KO and widespread neuronal KO of PP1β is lethal.

Secondly, we utilize the allocentric object-place assay to evaluate spatial learning and memory, since assays such as conventional novel-object recognition are not sensitive to lesions or perturbations of the hippocampus (Langston and Wood, 2009). While the use of our CA1-specific KO line aids in the interpretation of our results as the consequences of postsynaptic changes, it limits the utility of many behavioral assays of learning and memory. While the allocentric object-place assay was able to reveal the deficit in CA1 function seen in our PP1β cKO mice, it was unable to reveal any change in PP1α/γ cKO mice. It is unclear how sensitive this assay is to enhanced CA1 plasticity, specifically a decrease in the LTP threshold, and if modifications to the assay would improve its sensitivity, such as increasing the delay between the sample and test sessions. Regardless, we would expect that widespread neuronal KO of PP1α/γ would enhance spatial learning and memory in other behavioral assays.

## Acknowledgments

This work was supported by the National Institutes of Health (NIH) NIH F30 MH122046 to K.F., NIH DA10044 to A.C.N., R01 MH109719, R01 MH128279, National Science Foundation (NSF) IOS-1457336, University of Rochester Medical Center (URMC) Del Monte Neuroscience Institute Mangurian Foundation pilot program, and URMC Intellectual & Developmental Disabilities Research Center (IDDRC) Cell and Molecular Imaging (CMI) core pilot program to H.X.

## Author contributions

Conceptualization: KF, HX

Methodology: KF, CM, CM, DB, NW, HX

Investigation: KF, CM, AM, NW, YZ

Validation: AG

Visualization: KF

Resources: ACN, HX

Supervision: HX

Writing—original draft: KF, HX

Writing—review & editing: ACN

## Declaration of interests

The authors declare no competing interests.

## Figure Legends

**Fig. S1.**
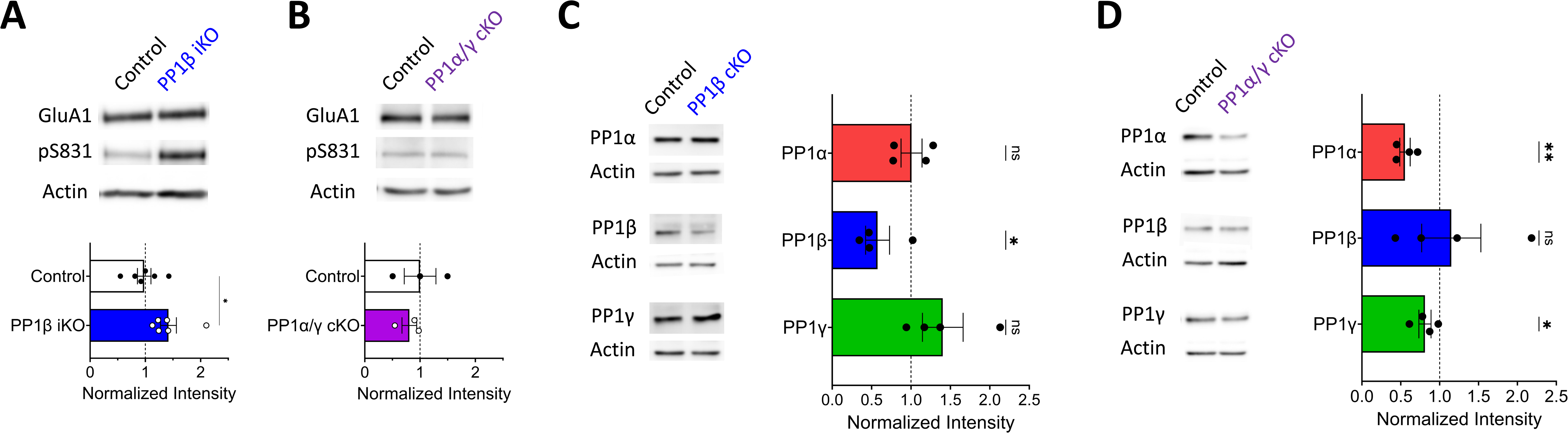
GluA1pS831 is a distinct target of PP1β and PP1β conditional knockout does not lead to PP1 isoform compensation at the protein level. (**A**) Western blot results from crude hippocampal synaptosomes from PP1β Thy1-CreER (iKO) mice and littermate controls. 6 mice per group. S831 GluA1 phosphorylation was significantly increased in PP1β iKO mice (two-tailed t-test: p<0.05). (**B**) Western blot results from CA1 microdissections from PP1α/γ cKO mice and littermate controls. 3 mice per group. There was no significant difference in GluA1 S831 phosphorylation (two-tailed t-test: p=0.57). **(C-D)** Western blot results from CA1 microdissections of (**C**) PP1β cKO and (**D**) PP1α/γ cKO mice and control littermates, showing a reduction in the knocked-out PP1 isoform at the protein level without a significant compensatory increase of the remaining PP1 isoform(s). 4 mice per group (one sample t-test: *p<0.1,**p<0.01).

**Fig. S2.**
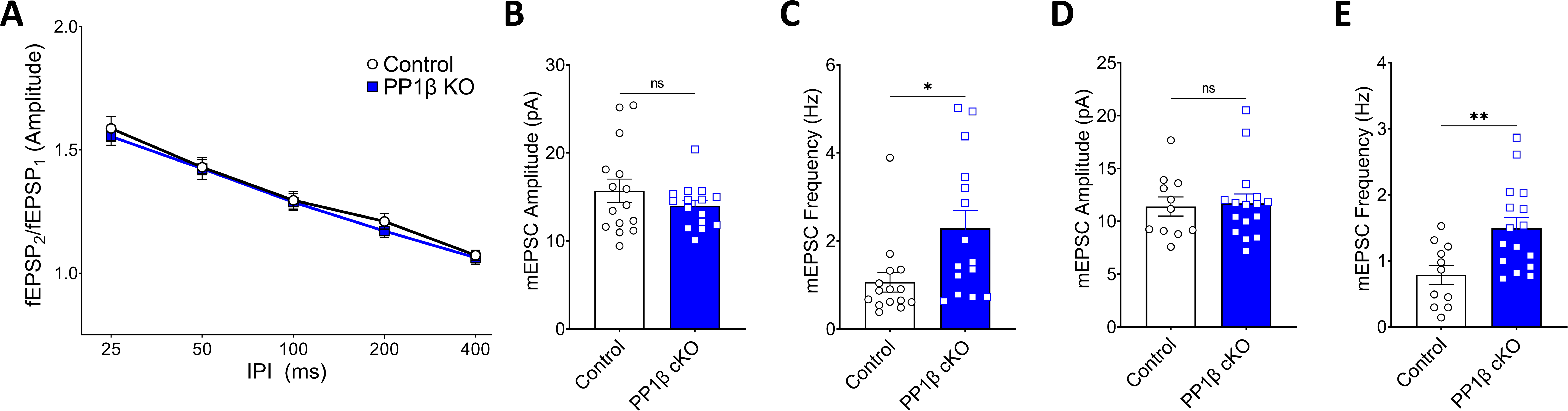
PP1β cKO mEPSC effects are independent of neurotransmitter release and acute hippocampal slice preparation methodology. (**A**) PP1β cKO has no effect on paired-pulse facilitation (two-way ANOVA: p=0.39, N=3 (control) and 3 (cKO) mice, n=10 and 10 slices). (**B-C**) mEPSC results from slices prepared in NMDG-based cutting solution. PP1β cKO has no effect on mEPSC amplitude (two-tailed t-test: p=0.25, N=3 and 3, n=15 and 15 cells) but increases mEPSC frequency (two-tailed t-test: p<0.05, N=3 and 3, n=15 and 15). (**D-E**) mEPSC results from slices prepared in sucrose-based cutting solution. PP1β cKO has no effect on mEPSC amplitude (two-tailed t-test: p=0.82, N=3 and 3, n=11 and 16) but increases mEPSC frequency (two-tailed t-test: p<0.01, N=3 and 3, n=11 and 16).

## STAR Methods

### Key resources table

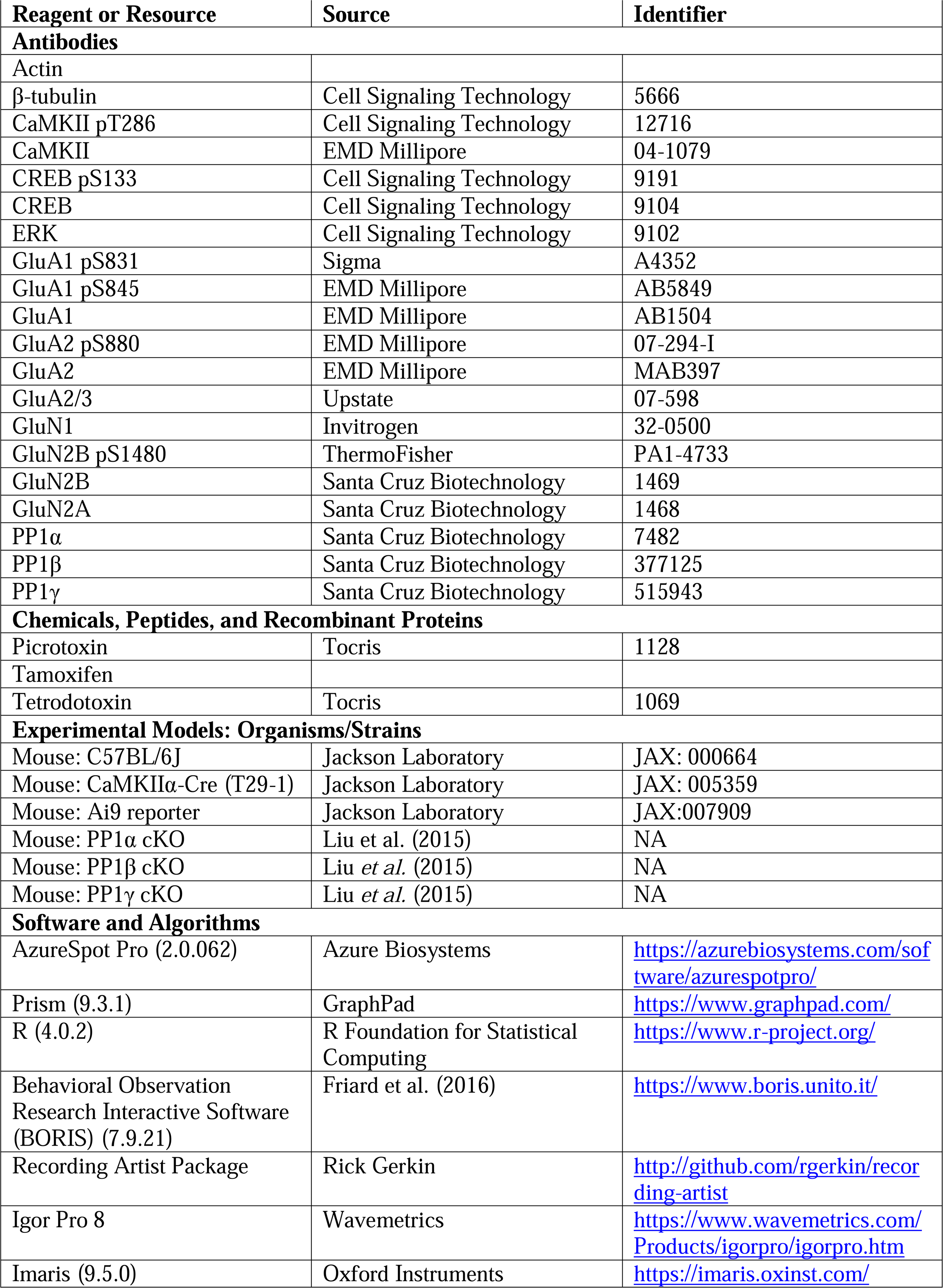

### Mice

CaMKIIα-Cre (T29-1; JAX 005359) and Thy1-CreER (JAX 012708) were purchased from Jackson laboratory. Floxed PP1 isoform mice were generated as described previously (Liu *et al*., 2015). Ai9 mice (JAX 007909) were a gift from A. Majewska at the University of Rochester. Mice were maintained by breeding with wild-type C57BL/6J (JAX 000664). Animals were group-housed (2-5 mice per cage) on a 12-hour light/dark cycle with food and water ad libitum. For all experiments, control littermates (PP1(fl/fl)) were compared to KO littermates (PP1(fl/fl); CaMKIIα-Cre; also called cKO). Thy1-CreER; PP1β(fl/fl) mice and control littermates were injected with tamoxifen (0.66mg/kg) starting at P21 for three days in a row (also called iKO). The mice were sacrificed at 3 weeks following injection. All animal procedures were conducted in accordance with the National Institutes of Health Guide for the Care and Use of Laboratory Animals and with protocols approved by the University of Rochester Institutional Animal Care and Use Committee.

### Acute hippocampal slice preparation

Acute hippocampal slices were prepared from 1-1.5 month-old mice of both sexes as described previously (Foley *et al*., 2022; Foley et al., 2021b). After brief isoflurane anesthetization and decapitation, the brain was rapidly dissected and sectioned in an ice-cold high sucrose solution containing, in mM: 215 sucrose, 2.5 KCl, 26 NaHCO_3_, 1.6 NaH_2_PO_4_, 20 D-glucose, 1 CaCl_2_, 4 MgCl_2_, and 4 MgSO_4_. 400 μm thick coronal (field recordings) or 300 μm thick transverse (whole-cell recordings) slices were sectioned on a Leica VT1200S vibratome with carbon steel blades (Electron Microscopy Sciences). After sectioning, slices were recovered in artificial cerebrospinal fluid (ACSF) at 34°C for 30 min, followed by an additional 1-hour incubation at room temperature (RT). ACSF contained, in mM: 124 NaCl, 2.5 KCl, 1.0 NaH_2_PO_4_, 26 NaHCO_3_, 10 D-glucose, 1.3 MgSO_4_, and 2.5 CaCl_2_ for field recordings; and 124 NaCl, 2.5 KCl, 1.2 NaH_2_PO_4_, 24 NaHCO_3_, 5 HEPES, 12.5 D-glucose, 2 MgSO_4_, and 2 CaCl_2_ for whole-cell recordings. For a subset of mEPSC recordings (Figure S2B,C), slices were prepared in NMDG-based ACSF containing, in mM: 92 NMDG, 2.5 KCl, 1.2 NaH_2_PO_4_, 30 NaHCO_3_, 20 HEPES, 25 D-glucose, 5 Na ascorbate, 2 thiourea, 3 Na pyruvate, 10 MgSO_4_, and 0.5 CaCl_2_. Solutions were continuously aerated with carbogen (95% O_2_, 5% CO_2_).

### Electrophysiology

Field recordings in acute hippocampal slices were performed at Sch-CA1 synapses in RT ACSF at a flow rate of 2-3 ml/min. Recording (∼1 MΩ) and stimulating (<1MΩ) electrodes were pulled from borosilicate glass and filled with ACSF. The recording electrode was placed in CA1 stratum radiatum, 200 μm medial to the stimulating electrode, with both electrodes approximately 100 μm below the slice surface. Responses were elicited every 15 sec. Input-output (IO) curves were measured by comparing the fiber volley amplitude to the fEPSP slope. Paired-pulse facilitation (PPF) ratios were measured by comparing the second fEPSP amplitude to the first. Long-term potentiation (LTP) was induced via 100 pulses, or 25 pulses (sub-threshold LTP), at 100 Hz after at least 10 min stable baseline recording.

Whole-cell recordings were performed on CA1 pyramidal neurons in ACSF held at 28°C (LTP; minimal stimulation) or 32°C (mEPSCs). Picrotoxin (50μM) was included in the ACSF for LTP and minimal stimulation, and picrotoxin (100μM) and tetrodotoxin (1μM) for mESPC recordings. 3-5 MΩ recording electrodes were pulled from borosilicate glass and filled with, in mM: 140 CsMeSO_3_, 8 CsCl, 7 Na_2_ phosphocreatine, 0.25 EGTA, 10 HEPES, 0.1 spermine, 2 Mg ATP, 0.3 Na GTP, at 7.32-7.36 pH and 294-298 mOsm for LTP and minimal stimulation, and 115 CsMeSO_3_, 20 CsCl, 10 Na_2_ phosphocreatine, 0.6 EGTA, 10 HEPES, 2.5 MgCl_2_, 5 QX-314-Br, 4 Mg ATP, 0.3 Na GTP, at 7.32-7.36 pH and 294-298 mOsm for mEPSC recordings. Stimulation electrodes were filled with ACSF. Responses were elicited every 10 sec and access and input resistance were continuously monitored. Cells were excluded from analysis if access resistance varied by more than 25%. CA1 pyramidal cells were identified via location (stratum pyramidale) and morphology under differential interference contrast microscopy with a 60x water-immersion objective on an Olympus BX51WI. For whole-cell LTP experiments, cells were depolarized to 0 mV during stimulation with 100 pulses at 100 Hz. For minimal stimulation experiments, stimulation intensity was reduced until ∼50% of stimulations failed to elicit an AMPAR-mediated response at a holding potential of -70 mV. We then measured the composite AMPAR- and NMDAR-mediated response by holding the cell at +40 mV. The percentage of silent synapses was calculated using the equation 1-ln(F_-70_)/ln(F_+40_), where F_-70_ is the failure rate at -70 mV holding potential and F_+40_ is the failure rate at +40 mV holding potential (Isaac et al., 1995; Liao et al., 1995). The stochastic nature of the release probability creates variability in the failure rate which can lead to mathematically negative values in the calculation of silent synapses (Favaro et al., 2018). Nevertheless, we excluded four PP1β cKO neurons which had calculated silent synapses proportions of <-0.15. No PP1β control neurons were excluded from the minimal stimulation analysis. For biocytin-fill experiments, neurons were held in whole-cell configuration for at least 10 min.

For all electrophysiology experiments, stimulation intensity was controlled by an ISO-Flex stimulus isolator (AMPI) and timing controlled by a Master-8 pulse stimulator (AMPI). Data were collected with a MultiClamp 700 series amplifier (Axon Instruments), PCI-6221 data acquisition device (National Instruments), and Igor Pro 7 (Wavemetrics) with a customized software package (Recording Artist, http://github.com/rgerkin/recording-artist).

### Spine density and morphology

Biocytin-filled neurons were fixed in 4% paraformaldehyde (PFA) for 24-48 hours and stained with conjugated streptavidin (Vector Laboratories) for 48 hours. Slices were then mounted with Prolong Diamond Antifade (Invitrogen) mounting media and allowed to cure for at least 24 hours. Confocal images of secondary dendrites within 200 μm of the cell body were acquired on an Olympus FV1000MP microscope system controlled by Fluoview software using a 60x oil immersion objective (Olympus UPlanSApo, 1.35 NA). Images were acquired at a resolution of 1024 x 1024 pixels at 3x zoom. For quantification of spine density and morphology, z-stacks of biocytin-filled neurons were deconvolved and reconstructed in 3D in Imaris (v9.3, Bitplane). Dendrites were reconstructed using semi-automated filament tracing in autopath mode. Spines were then identified using automatic seed points and manually verified. Spine identification parameters were set at 3.5 μm maximum spine length, 0.138 μm minimum spine head diameter, and branched spines were allowed. Spine morphology was classified in Imaris using the classify spines wizard algorithm. The general criteria for classification were: (i) Stubby spines have a large head without an appreciable neck; (ii) Mushroom spines have a large head greater than the width of the neck; (iii) Thin spines have a large head greater than the width of the neck, but smaller than mushroom spines; and (iv) Filopodia are long and lack an appreciable spine head. These criteria were achieved algorithmically with the following parameters in order, in μm: (i) Stubby: min(neck_width_) > 0.3 & length(spine) < 0.5 & max(head_width_) < max(neck_width_) | neck not detected; (ii) Mushroom: max(head_width_) > min(neck_width_)*2 & neck detected | max(head_width_) > 0.38; (iii) Thin: length(neck) > mean(neck_width_) & max(head_width_) < length(neck) | max(head_width_) > 0.14 ; and (iv) Filopodia: remaining spines. Note that spine morphology classification in Imaris is obligatorily ordered, such that spines that are classified as Stubby cannot be classified as Mushroom, and so on. Spine morphology was manually verified.

### SDS-PAGE and immunoblotting

CA1 microdissection was performed in ice-cold PBS after rapid extraction of the brain from 1-1.5 month-old mice anesthetized with isoflurane. Dissected tissue was frozen in liquid nitrogen and stored at -80°C until lysate preparation. Total protein lysates were prepared by homogenizing samples in RIPA buffer, containing 50 mM Tris-HCl pH 8.0, 1% NP-40, 0.5% Na deoxycholate, 0.1% SDS, 150 mM NaCl, 1 mM EDTA, 10 mM NaF, protease inhibitors (Pierce, A32965), and phosphatase inhibitor cocktails (Santa Cruz, 45055). Samples were agitated for 40 min at 4°C and centrifuged at 6,000 g for 20 min. The soluble fraction was used for protein concentration quantification (Pierce BCA) and then boiled in Laemmli buffer. Equal amounts of protein per sample were loaded and resolved by SDS-PAGE on 8-15% polyacrylamide gels and transferred to polyvinylidene fluoride membrane. Membranes were blocked with 5% bovine serum albumin or 5% non-fat dried milk in Tris-buffered saline with 0.1% Tween 20 (TBS-T) and washed with TBS-T. After primary antibody incubation (4°C), washing, secondary antibody incubation (RT), and final washing, immunoreactive signals were visualized by chemiluminescence on an Azure Biosystems Imaging System. Densitometry analysis was performed on AzureSpot Pro software (Azure Biosystems). The following antibodies were used: β-tubulin (Cell Signaling Technology (CST), 5666), CaMKII pT286 (CST, 12716), CaMKII (Millipore, 04-1079), CREB pS133 (CST, 9191), CREB (CST, 9104), ERK (CST, 9102), GluA1 pS831 (Sigma, A4352), GluA1 pS845 (Millipore, AB5849), GluA1 (Millipore, AB1504), GluA2 pS880 (Millipore, 07-294-I), GluA2 (Millipore, MAB397), GluA2/3 (Upstate, 07-598), GluN1 (Invitrogen, 32-0500), GluN2B pS1480 (PA1-4733), GluN2B (Santa Cruz, 1469), GluN2A (Santa Cruz, 1468).

For sufficient protein levels for fractionation studies, whole hippocampi from Thy1-CreER mice and control littermates were used. Hippocampi were rapidly dissected from mice euthanized via CO2 inhalation. Tissue was flash frozen in liquid nitrogen and stored at -80°C until crude synaptosomal preparation (Lautz et al., 2019). Each hippocampus was homogenized on ice with 12 strokes of the glass-glass Dounce Homogenizer in 1mL of sucrose buffer containing 320 mM sucrose, 5 mM HEPES, protease inhibitors (Pierce, A32965), and phosphatase inhibitor cocktail (Santa Cruz, 45055). Lysates were centrifuged at 1,0000 x g for 10 min at 4°C to separate the nuclear pellet (P1) and post nuclear supernatant (S1). The S1 fraction was centrifuged at 12,000 x g for 15 min at 4°C to isolate the light membrane fraction and soluble enzymes (S2) and crude synaptosome and mitochondria (P2). The P2 fraction was resuspended and agitated for 15 min at 4°C in lysis buffer containing 150 mM NaCl, 50 mM Tris-HCl pH 9.0, 10 mM NaF, 1% Na deoxycholate, protease inhibitors, and phosphatase inhibitor cocktail. P2 was re-centrifuged at 12,000 x g for 20 min at 4°C. The soluble fraction was collected and used as the crude synaptosome fraction, and the insoluble pellet was stored at -80°C. Protein quantification was performed on the synaptosome fraction before preparing samples containing 40 ug protein in Laemmli buffer for western blotting.

### Behavioral procedures

Behavioral experiments were performed on 1-1.5 month-old male mice, using approximately equal numbers of control and KO littermates from each litter. Three to four litters were used per mouse line. All experiments were carried out following 2-3 days of handling (Gouveia and Hurst, 2013) in the light phase (8 am-12 pm). Testing chambers and objects were thoroughly cleaned with 70% ethanol after each experiment. Open-field testing was performed for 1 hour in a chamber equipped with infrared photocells (Opto-Varimex-5, Columbus Instruments). Mouse position was automatically detected via beam interruption.

The allocentric object-place assay was performed following standard novel-object recognition procedures (Langston and Wood, 2009; Leger et al., 2013). The outer walls of the testing chamber were covered with four different colors or patterns for spatial cues. Sessions were split into three phases of 10 min each: habituation, sample, and test, with 5-10 min between phases to allow for thorough cleaning of the chambers. Three copies of the objects (A_1,2,3_, B_1,2,3_) were used to avoid any lingering cues between phases. In the habituation phase, mice were placed into the chamber facing the southern wall and allowed to explore the chamber in the absence of any objects. In the sample phase, two different objects (A_1_-B_1_) were placed in the chamber and mice were again placed facing the southern wall. In the test phase, two copies of one of the two sampling phase objects (A_2_-A_3_; B_2_-B_3_) were placed in the chamber and mice were placed into the chamber facing the eastern or western wall. Therefore, in the test phase, one of the objects was presented in the same location as in the sample phase (familiar; e.g. A_2_ at A_1_’s position), whereas the other object occupied a different location than in the sample phase (novel; e.g. A_3_ at B_1_’s position). Testing phase object pair (A or B) and site of entry (eastern/western wall) were counterbalanced. The objects used were 50 mL conical tubes (inverted) and 70 mL culture flasks filled with white or black sand. Exploration was defined as the mouse being within 2.5 cm of the object and directing its nose at the object while sniffing or whisking. Sessions were monitored via overhead video recording and analyzed in BORIS (Friard *et al*., 2016).

### Statistical analyses

GraphPad Prism (9.3.1) was used for statistical analyses and numerical data visualization. Statistical significance between means was calculated using unpaired, two-tailed t-tests or ANOVAs. Two-way ANOVAs were used for IO and PPF comparisons. IO curves were also fit with least squares regression curves without weighting and the slopes of the regressions were compared via extra sum-of-squares F-tests. LTP magnitude was quantified as the change in fEPSP slope at 25-30 min post-induction compared to the 5 (whole-cell) or 10 (field) minute baseline. Tukey and Šidák post-hoc comparisons were performed for one- and two-way ANOVAs, respectively. The arithmetic mean and standard error of the mean are displayed in all figures unless otherwise noted.

## Notes

### Competing Interest Statement

The authors have declared no competing interest.

### Summary of Updates

(1) We have shown that GluA1pS831 increase in the synaptosome fraction in PP1beta iKO mice. (2) We showed that there is no obvious change of PP1alpha and PP1gamma protein levels in the PP1b KO mouse hippocampi. (3) More discussions on the interplay between the 3 PP1 isoforms.

